# Quantification of bioactive compounds in the juices of 21 American elderberry cultivars

**DOI:** 10.1101/2025.11.12.688025

**Authors:** Amanda Dwikarina, Michael C. Greenlief, Andrew L. Thomas, Chung-Ho Lin

## Abstract

American elderberry (*Sambucus nigra* subsp. *canadensis*) is recognized as a rich source of bioactive compounds with nutritional and therapeutic potential. However, comprehensive quantification of its metabolites remains analytically challenging due to the structural complexity of phenolics and matrix interferences in juice extracts. In this study, a targeted ultra-high performance liquid chromatography-tandem mass spectrometry (UHPLC–MS/MS) workflow was developed and validated to quantify 22 bioactive metabolites, including anthocyanins, flavonoids, and phenolic acids, in juices from 21 American elderberry cultivars. An optimized UHPLC– MS/MS system demonstrated higher sensitivity, selectivity, and throughput than conventional high-performance liquid chromatography-tandem mass spectrometry (HPLC–MS/MS), achieving limits of detection and quantification below 0.1 ng/mL for most analytes. Matrix effects were effectively mitigated through sample dilution, enabling accurate and reproducible quantification across complex juice matrices. Quantification of metabolites showed that accession 1199 exhibited high levels of cyanidin-based anthocyanins, suggesting its utility as a natural food color source. Overall, high-throughput UHPLC–MS/MS method provides a robust analytical framework for metabolomic profiling for cultivar selection and the development of elderberry-derived functional products.

## 1. Introduction

American elderberry (*Sambucus nigra* subsp. *canadensis*) has gained popularity in recent decades as a rich natural source of bioactive phytochemicals with nutritional and therapeutic potential. The chemical composition of elderberries varies substantially among cultivars and plant tissues, suggesting the influence of both genetic and environmental factors on specialized metabolite biosynthesis^1–3^. Classes of compounds, including flavonoids, anthocyanins, and polyphenolic acids, were known to contribute to the characteristic dark purple color of elderberry ^1,2,4^. The chemical diversity within these classes results from various glycosylation, acylation, and methylation patterns that alter solubility, stability, and bioavailability^5^. This structural complexity presents analytical challenges in terms of extraction, separation, and accurate quantification of metabolites.

Comprehensive chemical identification and quantification of plant-derived metabolites is known to be challenging when applied to complex matrices^6^. Traditional analytical practices, such as gas chromatography-mass spectrometry (GC-MS), liquid chromatography with UV detection (LC-UV), and single-quadrupole liquid chromatography-mass spectrometry (LC-MS), are limited by low throughput, limited specificity, and labor-intensive sample preparation. Gas chromatography-mass spectrometry is known to be effective in analyzing volatile and low-polarity compounds^6,7^. The method development process in GC–MS is generally straightforward and easily transferable between instruments due to standardized ionization techniques and reproducible chromatographic conditions. However, for GC-MS, evaporation and derivatization steps during the sample preparation stage significantly increased the time required for sample preparation^6^. On the other hand, LC-UV methods, though cost-effective and widely applied for quality control, lack the selectivity to distinguish co-eluting isomeric compounds with overlapping UV absorption spectra. The LC-UV is still one of the analytical chemistry techniques that many people use, which can capture a broader range of non-volatile compounds^8–10^.

The development of LC-MS methods has addressed many of the limitations associated with traditional analytical techniques. The LC-MS offers greater selectivity and was optimized for the detection of a broad range of compounds^6^. Recent advances in LC separation technology, coupled with tandem mass spectrometry (MS/MS), have significantly increased analytical sensitivity and selectivity^8,6^. Advancements in column technology, such as hydrophilic-lipophilic balanced (HLB), hydrophilic interaction liquid chromatography (HILIC), and high-efficiency C18+ stationary phases, have further improved the resolution and peak capacity of compounds^11^.

Matrix interferences remain a common challenge in LC–MS/MS analysis of complex biological and plant-derived samples. These matrix effects (MEs) can alter ionization efficiency and compromise quantification accuracy^12,13^. Incorporating a sample clean-up step, such as solid-phase extraction (SPE), during sample preparation can reduce MEs; however, recovery rates and potential compound losses must be carefully evaluated to ensure accurate quantification^14^. Such dependencies highlight the need for optimized, compound-specific extraction and quantification strategies in metabolomic workflows.

Despite these advancements, comparative analysis of American elderberry juices using an optimized high-throughput LC-MS/MS workflow remains limited. Previous studies have focused on a selected group of chemicals, such as cyanidin^1–3^. A robust, validated targeted metabolomics approach is therefore needed to quantify a broad range of bioactive compounds across diverse cultivars and to identify genotype-specific metabolic signatures that may inform breeding, product development, and the standardization of elderberry-based nutraceuticals.

This study aimed to develop and validate a streamlined LC-MS/MS workflow for quantifying major bioactive anthocyanins, flavonoids, and phenolic acids in American elderberry juices. Using both high-performance liquid chromatography-tandem mass spectrometry (HPLC-MS/MS) and ultra-high-performance liquid chromatography-tandem mass spectrometry (UHPLC-MS/MS), this study compared analytical performance, evaluated matrix effect, and quantified metabolites in 21 cultivars of American elderberry.

## 2. Materials and Methods

### 2.1. Chemical

Acetonitrile, and formic acid were purchased from Fisher Chemical (Fair lawn, NJ, USA) and were all LC/MS grade. Methanol for sample extraction were of HPLC-grade and purchased from Sigma-Aldrich (St. Louis, MO, USA). Chemical standards for standard calibration curve were purchased from Sigma-Aldrich (St. Louis, MO, USA) with purity >95%, except caffeic acid, catechin, p-coumaric acid, cyanidin 3,5-*O*-diglucoside, cyanidin 3-*O*-glucoside, epicatechin, gallic acid, neochlorogenic acid, and isorhamnetin 3-*O*-glucoside with purity >98%; kaempferol with purity >97%; myricetin with purity >96%; cyanidin 3-*O*-rutinoside, peonidin 3-*O*-glucoside and isoquercetin with purity >90%; ferulic acid was of certified reference standard grade; and isorhamnetin 3-rutinoside was purchased from Fisher Chemical (Fair lawn, NJ, USA) with purity >95%.

### 2.2. Plant materials

The elderberry juices samples analyzed in this study have been previously described in Dwikarina et al. (2024). American elderberry (*Sambucus nigra* subsp. *canadensis*) fruits from 18 propagated accessions and three established cultivars were harvested from plantings at the University of Missouri’s Southwest Research, Extension, and Education Center near Mt. Vernon, Missouri, USA. The fruits were harvested at peak ripeness, and the juice was pressed by hand. The juices were then filtered through a kitchen sieve, aliquoted into 50 mL polypropylene tubes, and were kept frozen at -20 °C until further analysis.

### 2.3. Sample preparation

One mL of raw juice was mixed with three mL of methanol in a glass vial. The mixture was then sonicated for 60 minutes using a sonicator (Fisher Scientific, Pittsburgh, PA, USA). The extract then centrifuged for 30 min at 5000 rpm with temperature set at 10 °C. One mL of juice extract (supernatant) was transferred to a 1.5 mL Eppendorf tubes. The extract was then filter using 25 mm 0.2µm PTFE filter (Acrodisc syringe filter). 100 µL of the juice extract was diluted with methanol to final dilution factor of 800. 1.5 mL of diluted juice extract was then transferred to the LC-MS /MS vial prior to injection to HPLC-MS/MS. 750 µL of the diluted juices extract was further diluted with methanol (1:2, v/v) then transferred to LCMS vial prior to injection to UHPLC-MS/MS.

### 2.4. HPLC-MS/MS Methods

The targeted metabolomic analysis of compound chlorogenic acid and neochlorogenic acid were performed on Waters Alliance 2695 Separation Module High Performance Liquid Chromatography system coupled with a Waters Acquity TQ triple quadrupole mass spectrometer (HPLC-MS/MS) consisting of a quaternary pump, autosampler and a column oven. The analytical column for HPLC was a Phenomenex (Torrance, CA, USA) Kinetex C18 100 Å (100 length x 4.6 mm internal diameter, 2.6 µm particle size) reversed-phase column. Sample injection volume was 30 µl. Column temperature was maintained at 40°C ± 5°C. A linear two-part mobile-phase gradient was used. Mobile phase A consisted of 100% acetonitrile, and mobile phase B consisted of 0.1% formic acid in water. The gradient conditions are: 0 - 0.3 min, 2% A; 0.3 - 7.27 min, 2-80% A (linear gradient); 7.27 – 7.37 min, 80-98% A (linear gradient); 7.37 – 9 min, 98% A, 9 – 10 min, 98-2% A (linear gradient), and the gradient was maintained at 2% A until 15 min. The flow rate was set at 0.5 mL/min.

To identify the chemical compound in the American elderberry sample, the MS/MS system was operated using electrospray ionization (ESI) in negative ionization mode with capillary voltage of kV. The ionization source was programmed at 150°C and the desolvation flow rate was programmed at 750 L h^-1^. The MS/MS system was operated in multi-reaction monitoring (MRM) mode with parameters as described in Table 1. The Retention time of neochlorogenic acid was around 5.71 min, while the retention time of chlorogenic acid was around 6.07 min.

**Table 1.** Optimized multiple reaction monitoring (MRM) conditions for the analysis of neochlorogenic acid and chlorogenic acid in HPLC-MS/MS.

| Compound name | MW (g/mol) | Ion mode | RT (min) | Q1 (*m/z*) | Q2 (*m/z*) | Cone (V) | Collision (eV) |
| --- | --- | --- | --- | --- | --- | --- | --- |
| Neochlorogenic acid | 354.10 | ES<sup>-</sup> | 5.71 | 352.95 | 191.3 | 40 | 20 |
| Chlorogenic acid | 354.10 | ES<sup>-</sup> | 6.07 | 352.95 | 191.3 | 40 | 20 |

\* Q1 = Molecular ion, Q2 = Product ion (quantifier)

The MS/MS system was optimized using collision optimization in the MRM mode. The transition ions for molecular and product ions identification, MRM, cone voltage, and collision energy were optimized by Empower 3 Autotune Wizard software package. Analytical data were processed using Waters Empower 3 software (Waters, USA).

### 2.5. UHPLC-MS/MS Method

The concentrations of American elderberry compounds were determined by a Waters Acquity Ultra-High-Performance Liquid Chromatography (Waters, Milford, MA, USA) coupled with a XEVO TQ-XS tandem mass spectrometer (UHPLC-MS/MS, Waters) controlled by MassLynx software Ver 4.2. The compounds were separated by a CORTECS® C18+ analytical column with µm particle size, 100 mm length x 2.1 mm internal diameter connected to CORTECS® UPLC C18+ VanGuard Pre-Column (1.6 µm particle size, 5 mm length x 2.1 mm internal diameter). Separation was achieved using a linear gradient of 0.01% formic acid in water (A) and 0.01% formic acid in 100% acetonitrile (B). The gradient conditions are: 0–0.2 min, 2% B; 0.2–1.89 min, 2–80% (linear gradient) B; 1.89-1.92 min, 80–98% (linear gradient) B; 1.92–3.61 min (linear gradient), 98% B; 3.61–6.77 min, 2% B with a flow rate of 0.4 ml/min. The column temperature was set at 40°C and the autosampler temperature was set at 10°C. The system was first conditioned with 50 % acetonitrile and 50% of 0.01% formic acid, and the column was equilibrated with 2% acetonitrile and 98% of 0.01% formic acid solution before injection. The injection volume is 2 µl. The mass analyzer Xevo-TQXS was equipped with an electrospray ionization (ESI) source and operated in positive and negative ion mode. The acquisition parameters for compounds were carried out in the multi-reaction monitoring mode (MRM). Two fragment ions were monitored for quantification of analytes and qualifier as trace ions for confirmation (Table 2). The ionization energy, multi-reaction monitoring (MRM) transition ions (precursor and product ions), capillary and cone voltage (CV), desolvation gas flow, and collision energy (CE) were optimized by Waters IntelliStart™ optimization software package. The data was processed, quantified, and reviewed by TargetLynx software Ver 4.2. The optimized ionization, collision energy, and ion selections were described as in Table 2.

**Table 2.** Optimized MRM conditions for the analysis of elderberry compounds in UHPLC-MS/MS.

| Compound Name | MW (g/mol) | Ion mode | RT (min) | Q1 (*m/z*) | Q2 (*m/z*) | Cone (V) | Collision (eV) | Q3 (*m/z*) | Cone (V) | Collision (eV) |
| --- | --- | --- | --- | --- | --- | --- | --- | --- | --- | --- |
| Cyanidin 3,5-*O*-diglucoside | 611.16 | ES<sup>+</sup> | 2.06 | 611.235 | 449.152 | 44 | 18 | 287.078 | 44 | 36 |
| Cyanidin 3-*O*-sophoroside | 611.16 | ES<sup>+</sup> | 2.07 | 611.235 | 136.928 | 50 | 70 | 287.078 | 50 | 28 |
| Cyanidin 3-*O*-sambubioside | 581.15 | ES<sup>+</sup> | 2.09 | 581.220 | 287.080 | 44 | 28 | 136.93 | 44 | 68 |
| Cyanidin 3-*O*-galactoside /<br>Cyanidin 3-*O*-glucoside | 449.11 | ES<sup>+</sup> | 2.10 | 449.185 | 287.085 | 34 | 26 | 136.995 | 34 | 54 |
| Cyanidin 3-*O*-rutinoside | 595.17 | ES<sup>+</sup> | 2.10 | 595.181 | 287.078 | 58 | 28 | 136.929 | 58 | 72 |
| Pelargonidin 3-*O*-glucoside | 433.40 | ES<sup>+</sup> | 2.14 | 432.900 | 271.104 | 32 | 30 | 120.995 | 32 | 54 |
| 3-4, Dihydroxybenzoic acid | 154.03 | ES<sup>-</sup> | 2.15 | 152.998 | 108.856 | 12 | 14 | 80.857 | 12 | 22 |
| Peonidin 3-*O*-glucoside | 463.40 | ES<sup>+</sup> | 2.16 | 463.202 | 301.122 | 28 | 18 | 286.034 | 28 | 44 |
| Catechin | 290.08 | ES<sup>-</sup> | 2.22 | 289.110 | 245.037 | 40 | 16 | 202.876 | 40 | 20 |
| Epicatechin | 290.27 | ES<sup>-</sup> | 2.24 | 289.110 | 245.037 | 46 | 16 | 108.897 | 46 | 24 |
| Caffeic acid | 180.40 | ES<sup>+</sup> | 2.30 | 181.064 | 88.986 | 18 | 28 | 162.998 | 18 | 8 |
| Delphinidin 3-*O*-rutinoside | 611.16 | ES<sup>+</sup> | 2.30 | 611.171 | 303.016 | 50 | 30 | 228.991 | 50 | 70 |
| Rutin (Quercetin-3-rutinoside) | 610.50 | ES<sup>-</sup> | 2.30 | 609.341 | 300.145 | 78 | 36 | 271.007 | 78 | 64 |
| Myricetin | 318.23 | ES<sup>-</sup> | 2.33 | 317.070 | 150.000 | 6 | 28 | 178.86 | 6 | 20 |
| Isoquercetin / Hyperoside | 464.40 | ES<sup>-</sup> | 2.34 | 463.177 | 300.165 | 60 | 26 | 270.964 | 60 | 44 |
| Kaempferol 3-*O*-rutinoside | 594.50 | ES<sup>+</sup> | 2.35 | 595.277 | 287.065 | 30 | 20 | 449.191 | 30 | 12 |
| Isorhamnetin 3-rutinoside | 624.50 | ES<sup>-</sup> | 2.36 | 623.330 | 314.857 | 74 | 32 | 270.95 | 74 | 56 |
| Kaempferol 3-glucoside | 448.40 | ES<sup>+</sup> | 2.39 | 449.177 | 287.082 | 26 | 12 | 153.002 | 26 | 48 |
| Isorhamnetin 3-*O*-glucoside | 478.40 | ES<sup>-</sup> | 2.40 | 477.177 | 314.061 | 68 | 28 | 242.907 | 68 | 44 |
| p-Coumaric acid | 164.16 | ES<sup>-</sup> | 2.41 | 163.000 | 118.941 | 16 | 14 | 92.834 | 16 | 26 |
| Ferulic acid | 194.18 | ES<sup>-</sup> | 2.45 | 193.020 | 133.898 | 10 | 16 | 177.893 | 10 | 12 |
| Luteolin | 286.24 | ES<sup>+</sup> | 2.61 | 287.080 | 134.989 | 42 | 32 | 153.016 | 42 | 28 |
| Quercetin | 302.23 | ES<sup>-</sup> | 2.62 | 301.070 | 150.876 | 48 | 22 | 178.859 | 48 | 20 |
| Naringenin | 272.25 | ES<sup>+</sup> | 2.72 | 273.090 | 153.010 | 46 | 20 | 147.024 | 46 | 18 |
| Kaempferol | 286.24 | ES<sup>+</sup> | 2.75 | 287.080 | 120.997 | 46 | 30 | 153.016 | 46 | 32 |
| Isorhamnetin | 316.26 | ES<sup>-</sup> | 2.79 | 315.100 | 300.000 | 56 | 22 | 150.89 | 56 | 28 |

\*Q1 = Molecular ion, Q2 = Product ion (quantifier), Q3 = Trace ion (qualifier)

### 2.6. Method validation

The optimized method was validated for linearity, limit of detection (LOD), and limit of quantification (LOQ), carry-over, and matrix effects. Linearity of each compound was assessed by injection of seven-point calibration curves ranging from 5 to 500 ng mL^-1^ for analysis in HPLC-MS/MS and from 0.1 to 100 ng mL^-1^ for analysis in UHPLC-MS/MS. LOD and LOQ values were calculated using the following equation:

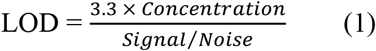

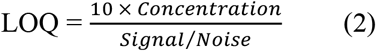

Carry-over was assessed by injecting five solvent-blank samples after the highest calibration standard concentration. To assess the matrix effect (ME) and recovery (RE) of compound in HPLC-MS/MS analysis^15^, a standard mixture solution (100 ng mL^-1^) was added to the 800-fold diluted elderberry juice extract and calculated as

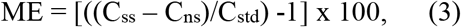

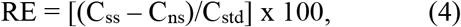

where C_ss_, C_ns_ and C_std_ represent the estimated concentration of the spiked sample, a non-spiked sample, and a spiked standard concentration in the solvent, respectively. To assess the matrix effect in UHPLC-MS/MS analysis, a standard mixture solution (10 ng mL^-1^) was added to the 1,600-fold diluted elderberry juice extract. The matrix effect and recovery rate of the compound were calculated using equations (3) and (4), respectively.

### 2.7. Statistical analysis

Quantification of analytes in American elderberry juices was performed in three analytical replicates. The results were expressed as mean ± SD. One-way ANOVA was performed using the statistical program R (ver. 4.5.1) to assess significant differences in analyte concentrations between cultivars. Statistical significance was defined at *p-*value of < 0.001. Multivariate analysis was performed using Metaboanlyst ver. 6.0 (https://www.metaboanalyst.ca/).

## 3. Results

The analytes in this study were analyzed using HPLC–MS/MS and UHPLC–MS/MS platforms. Based on the calculated limits of detection (LOD) and quantification (LOQ), the UHPLC–MS/MS system demonstrated better sensitivity and overall performance (Table 3). However, chlorogenic acid (CGA) and neochlorogenic acid (NCGA) were not detected by UHPLC–MS/MS, likely due to the differences in column chemistry and mobile-phase composition. Furthermore, evidence of matrix effect (ME) was observed on several analytes when analyzed by HPLC-MS/MS (data not shown). To minimize this effect, samples were further diluted. Because dilution reduces analyte concentration, the use of a more sensitive analytical platform was essential to maintain the detection capability. This approach effectively reduced MEs to below 20% for most analytes, except for quercetin (Table 4-5). Nevertheless, all quantitative results were corrected by the estimated MEs value. All compounds except CGA and NCGA were quantified using UHPLC– MS/MS, while these two phenolic acids were analyzed using HPLC–MS/MS.

**Table 3.** Calculated LOD and LOQ of selected analytes as measured in HPLC-MS/MS and UHPLC-MS/MS.

| Compound name | HPLC-MS/MS |  | UHPLC-MS/MS |  |
| --- | --- | --- | --- | --- |
|  | LOD* | LOQ* | LOD* | LOQ* |
| 3-4, Dihydroxybenzoic acid | 1.572 | 5.188 | 0.850 | 2.806 |
| Caffeic acid | 0.474 | 1.564 | 0.319 | 1.054 |
| p-Coumaric acid | 1.646 | 5.430 | 0.150 | 0.496 |
| Quercetin | 0.896 | 2.957 | 0.083 | 0.273 |
| Kaempferol | 0.217 | 0.717 | 0.009 | 0.029 |
| Naringenin | 0.098 | 0.324 | 0.012 | 0.038 |
| Cyanidin 3,5-O-diglucoside | 3.914 | 12.917 | 0.038 | 0.125 |

\*LOD and LOQ concentrations are in ng/mL

**Table 4.**
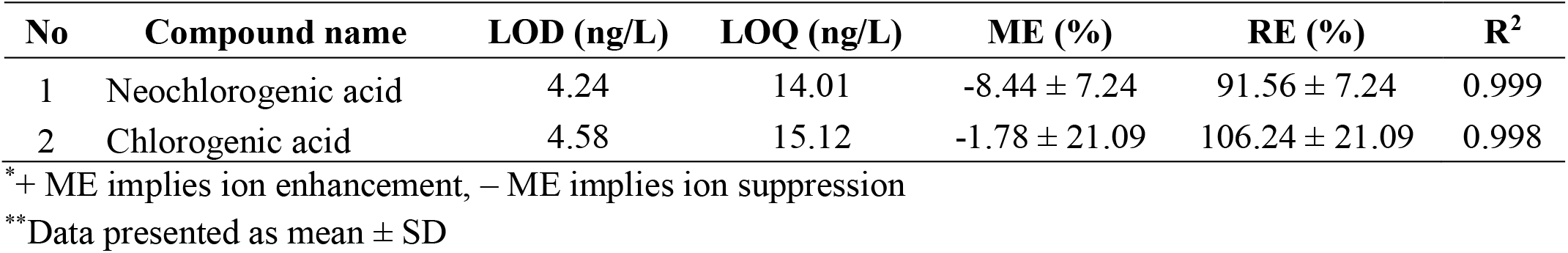
Calculated LOD, LOQ, and mean recovery of matrix effect of analytes for spike concentrations of 100 ng mL^-1^ measured by HPLC-MS/MS.

| No | Compound name | LOD (ng/L) | LOQ (ng/L) | ME (%) | RE (%) | R<sup>2</sup> |
| --- | --- | --- | --- | --- | --- | --- |
| 1 | Neochlorogenic acid | 4.24 | 14.01 | -8.44 ± 7.24 | 91.56 ± 7.24 | 0.999 |
| 2 | Chlorogenic acid | 4.58 | 15.12 | -1.78 ± 21.09 | 106.24 ± 21.09 | 0.998 |

\*+ ME implies ion enhancement, – ME implies ion suppression
\*\*Data presented as mean ± SD

Method optimization for each analyte was performed with Waters IntelliStart™ software to generate product ions and optimize multiple reaction monitoring (MRM) parameters (Table 1). The method optimized for CGA and NCGA was performed using the AutoTune Wizard package in Empower3 software, and the optimized method parameters are presented in Table 2. An example of ion spectra generated for cyanidin 3,5-*O*-diglucoside by Waters IntelliStart™ software is shown in Figure 1. The molecular ion *m/z* 611.235 was detected in positive ionization mode, corresponding to the protonated ion [M^+^] of cyanidin 3,5-*O*-diglucoside. Fragmentation of this precursor ion produced product ions at *m/z* 449.152 and *m/z* 287.078, corresponding to the sequential loss of one and two glucose moieties, respectively. The fragment *m/z* 287.078 corresponds to the aglycone cyanidin, which is a characteristic of cyanidin-based anthocyanins. This fragmentation pattern is consistent with previous reports that characterized cyanidin derivatives in dried red hybrid-tea rose petals using high-mass-accuracy, multi-dimensional fragmentation analysis by liquid chromatography–electrospray ionization–ion trap–time of flight mass spectrometry (LC–ESI–IT–TOF–MS)^16^.

**Figure 1.**
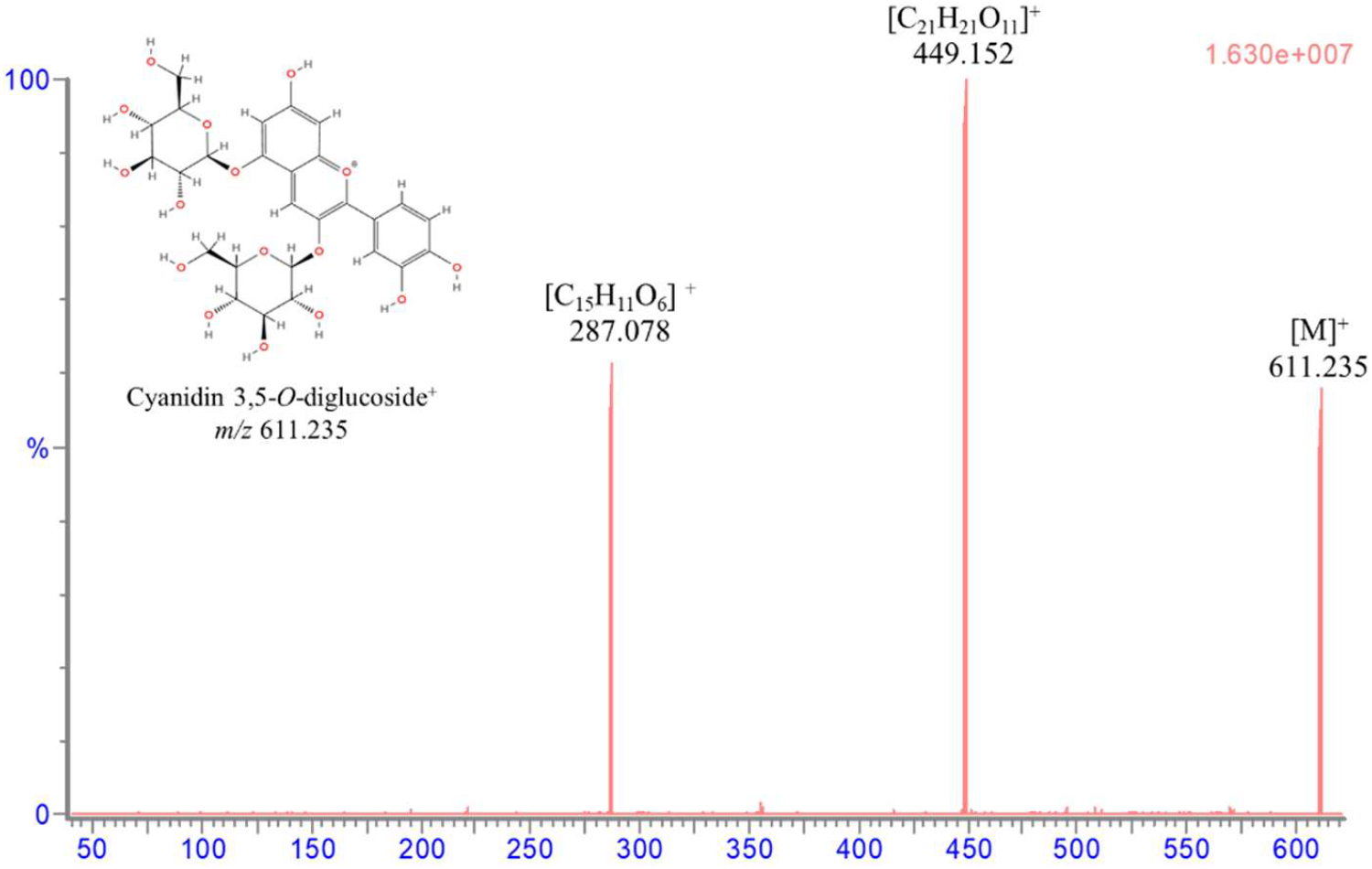
Ion spectra generated by the Waters Intellistart software package of the molecular ion *m/z* 611.235 and product ions at *m/*z 449.152 and *m/z* 287.078 for the analysis of cyanidin 3,5-*O*-diglucoside in positive ion mode.

Two cyanidin isomers, cyanidin 3-*O*-galactoside and cyanidin 3-*O*-glucoside, could not be chromatographically separated due to the identical retention time and fragmentation patterns; hence, the quantification of these analytes should be considered as the sum of two isomeric compounds. Similarly, quercetin derivatives, isoquercetin and hyperoside, which differ only in the attached sugar moiety, were not chromatographically resolved under the current analytical conditions, and their quantification was likewise reported as the sum of the two isomeric compounds.

For quantification of analytes in American elderberry juices, seven-point calibration curves ranging from 5 to 500 ng mL^-1^ for analysis in HPLC-MS/MS and from 0.1 to 100 ng mL^-1^ for analysis in UHPLC-MS/MS were used. Standard calibration curves showed good linear correlations (*R*^*2*^ values >0.998) between integrated peak areas and known compound concentrations. An example of a standard calibration curve is shown in Figure 2A with an integrated peak at RT 2.05 min for the quantification of compound cyanidin 3,5-*O*-diglucoside.

**Figure 2.**
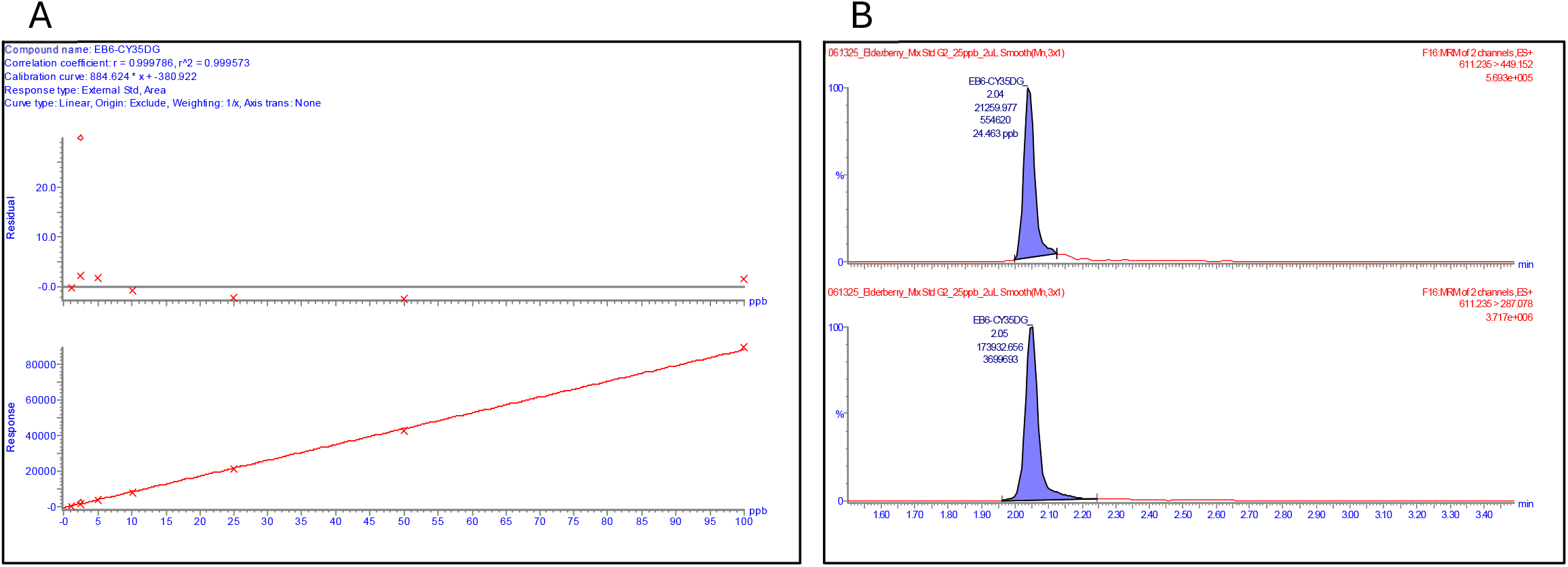
Analysis of compound cyanidin 3,5-*O*-diglucoside. A. Standard calibration curve (R^2^ = 0.9996). B. Ion chromatogram of compound at RT 2.05 min.

An attempt to quantify myricetin in American elderberry juices was carried out using both HPLC-MS/MS and UHPLC-MS/MS systems; however, the compound exhibited poor stability and was found to be degraded relatively quickly, even in stock standard solutions. This instability is consistent with a previous report describing the susceptibility of myricetin and related flavonols to oxidation under ambient conditions^17^. Therefore, the quantification of myricetin was excluded from the analysis.

Juices from 21 American elderberry cultivars were analyzed and quantified for their concentrations of three major classes of compounds: anthocyanins, phenolic acids, and flavonoids. The results were presented in Figures 3-6 for the quantification of anthocyanins, phenolic acids, and flavonoids, respectively. The results showed that cyanidin-3-O-rutinoside was detected only in accession 1199, with a concentration of 2.54 ± 0.12 mg/L. Cyanidin-3-*O*-sophoroside was found to be the highest in accession 1896. Cyanidin-3-*O*-glucoside was detected in 5 accessions, with the highest concentration found in accession 1199, followed by accession 2083. Phenolic acid compounds were found to be the most abundant in accession 1196, although CGA and NCGA were not detected in this accession. In contrast, the flavonoid compound isorhamnetin 3-rutinoside was found to be the highest in cultivar Ozark.

**Figure 3.**
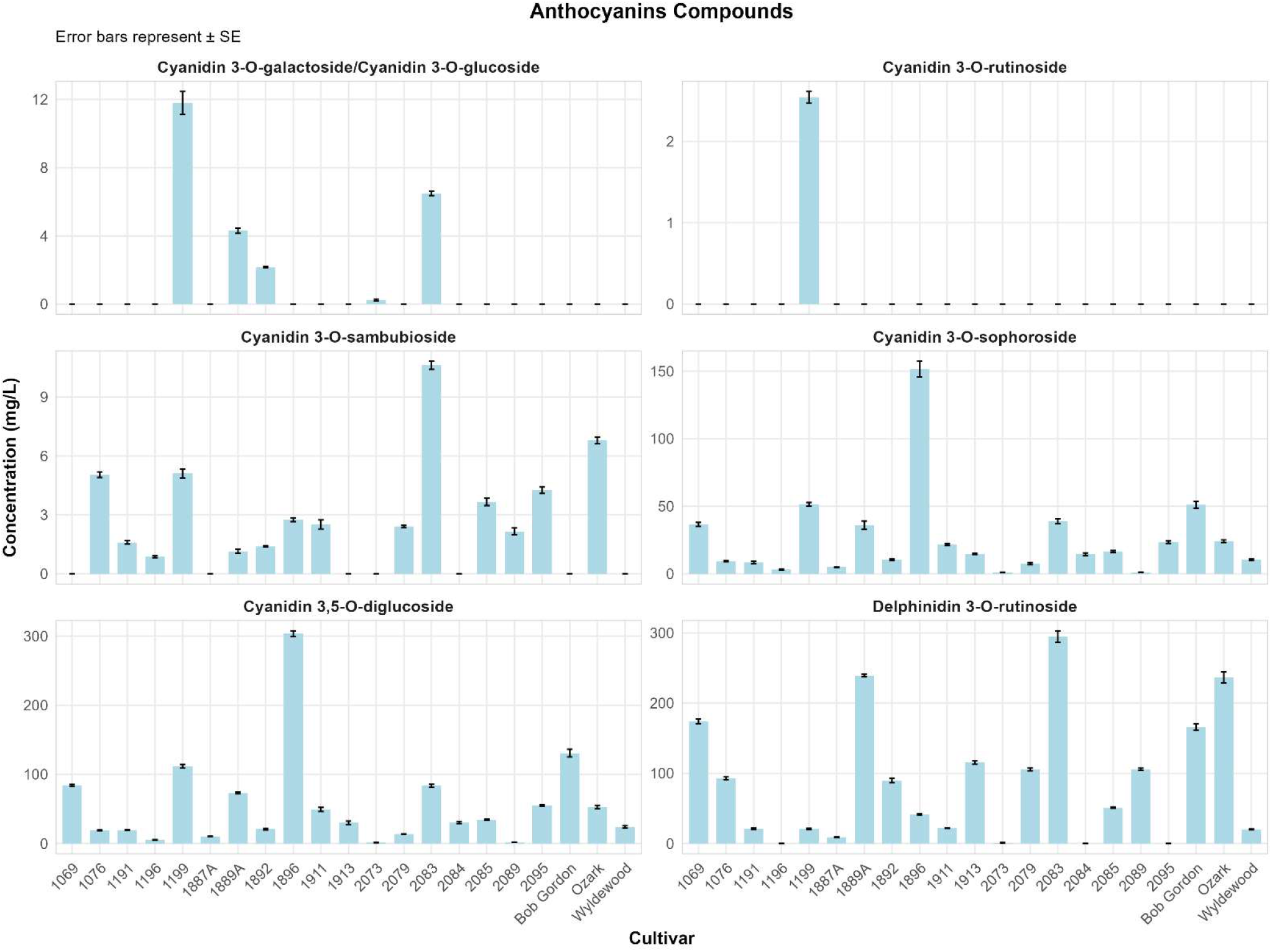
Mean concentration (mg/L) of anthocyanin compounds quantified in the juice of American elderberry cultivars measured by UHPLC-MS/MS. Results were presented as mean value ± standard error (n=3).

**Figure 4.**
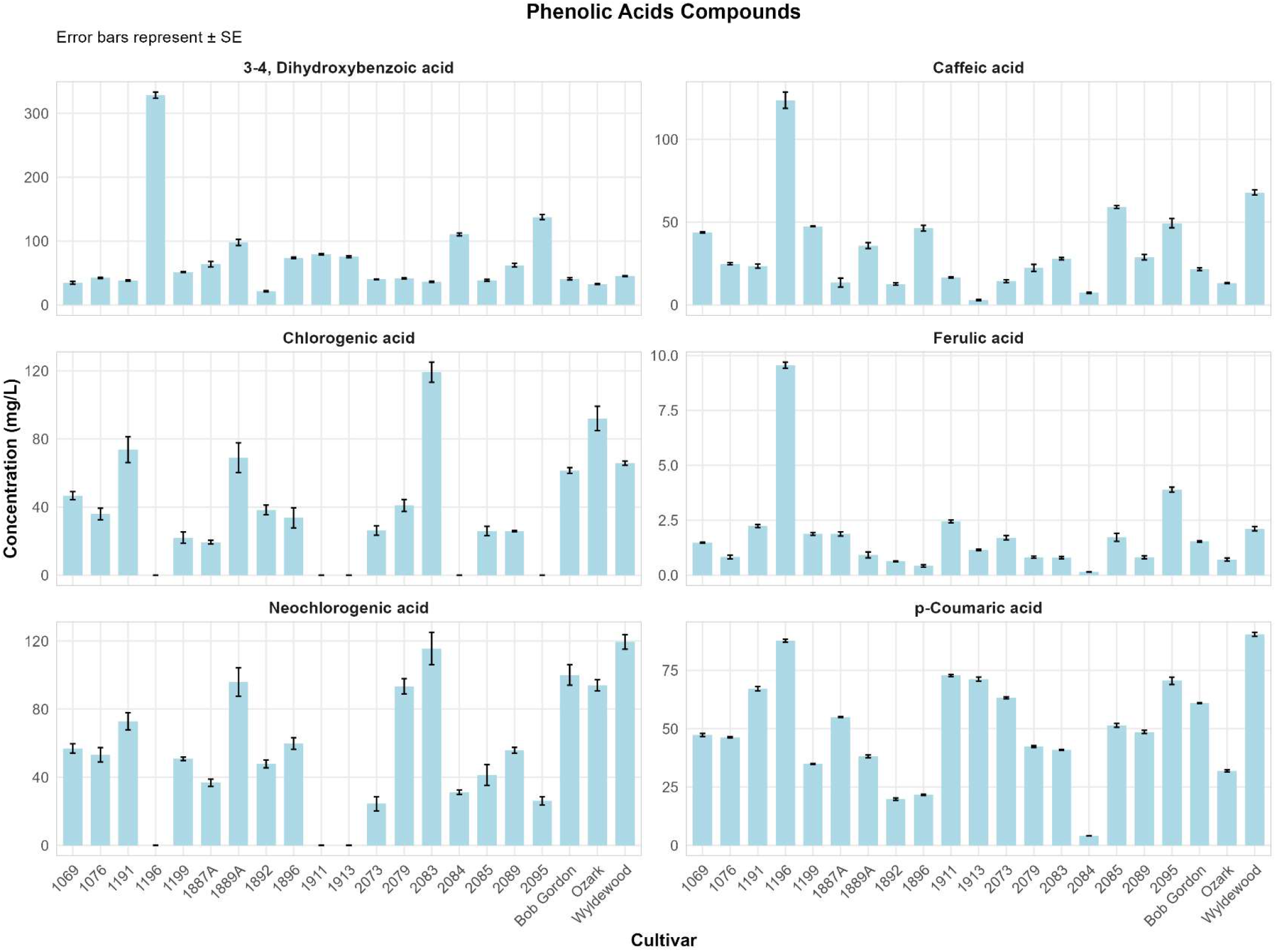
Mean concentration (mg/L) of phenolic acid compounds quantified in the juice of American elderberry cultivars measured by UHPLC-MS/MS and HPLC-MS/MS. Results were presented as mean value ± standard error (n=3).

**Figure 5.**
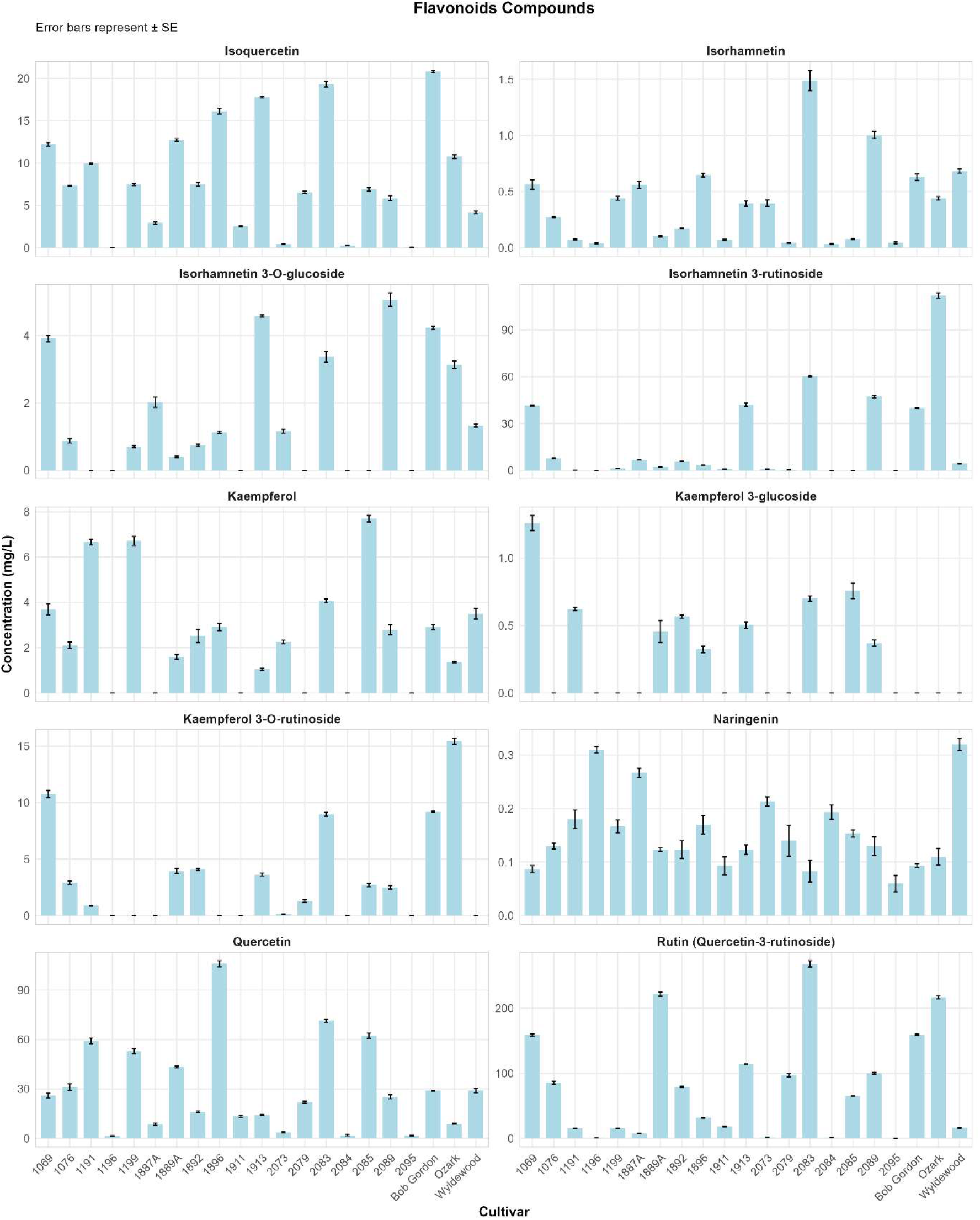
Mean concentration (mg/L) of flavonoid compounds quantified in the juice of American elderberry cultivars measured by UHPLC-MS/MS. Results were presented as mean value ± standard error (n=3).

**Figure 6.**
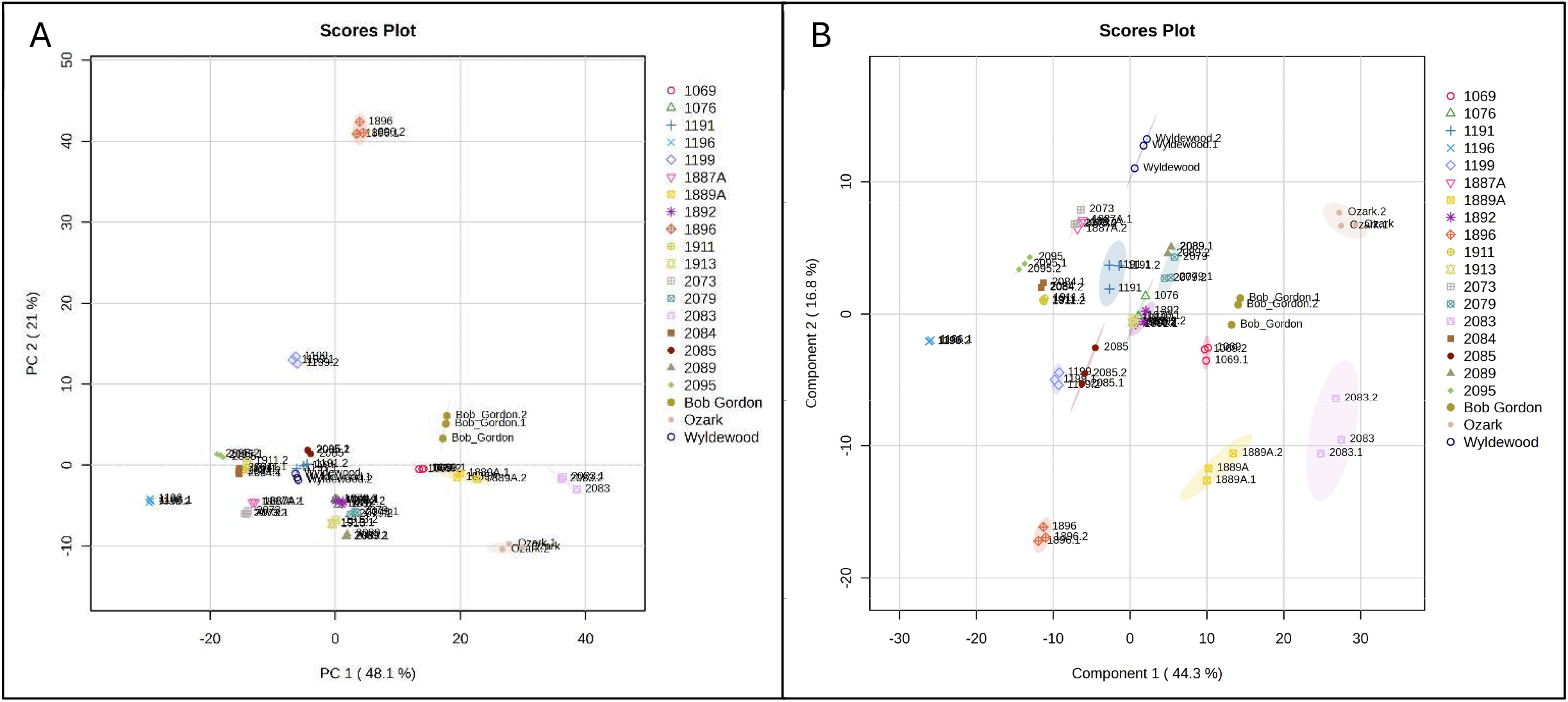
Multivariate analysis of compounds in various cultivars of American elderberry juices. **A.** PCA scores plot and **B**. PLS-DA scores plot of compounds in juices of 21 American elderberry cultivars revealed differences in the metabolomic profiles between cultivars. Circles with the same color represent analytical replicates (n=3). The colored ellipses indicate 95% confidence regions of each group.

An unsupervised multivariate analysis, principal component analysis (PCA), was performed to explore overall patterns in the metabolite profile of American elderberry (Figure 6A). The PCA results showed separation among cultivars, indicating differential metabolic profiles between groups. Two principal components (PCs) explained 69.1% of the total variability, with the first principal component (PC1) explaining 48.1% of the data variability and the second PC (PC2) accounting for 21% of the data variability. A supervised model, partial least-squares discriminant analysis (PLS-DA)^18^, was applied to further resolve group-specific differences (Figure 6B). PLS-DA model revealed significant discrimination among American elderberry cultivars, with two components explaining 61.1% of the total variance (PC1: 44.3%; PC2: 16.8%). Notably, accession 1896 clustered distinctly from the other cultivars, suggesting substantial differences in its metabolite composition. Cultivar Ozark also clustered separately from the other cultivars, indicating a distinct metabolite composition. This observation is consistent with previous studies employing untargeted metabolomic approaches, which similarly reported unique chemical profiles for Ozark compared to other American elderberry genotypes^19,20^.

## 4. Discussion

This study aimed to establish a rapid, high-throughput method for quantifying classes of bioactive metabolites in American elderberry juices. Both HPLC-MS/MS and UHPLC-MS/MS systems were evaluated for their analytical performance. The UHPLC-MS/MS setup demonstrated a higher sensitivity with better chromatographic resolution and shorter run times compared to the HPLC-MS/MS system. The smaller particle size in the UHPLC column improved the column efficiency, resulting in lower baseline noise and higher signal-to-noise ratios. The sensitivity and selectivity of UHPLC-MS/MS at low detection levels enable precise quantification of analytes in complex matrices with minimal sample preparation.

Complex sample matrices are known to interfere with analytical measurements. The matrix effect (ME) refers to the direct or indirect alteration of an analyte’s response caused by co-eluting, unintended components present in the sample. These interferences can either suppress or enhance ionization efficiency in the mass spectrometer, leading to inaccurate quantification if not properly monitored^12,13,21^. In this study, a 1600-fold dilution of the samples minimized the matrix effect, as shown in Table 5 with range of recovery rates of 82.74% ± 3.19 to 139.1% ± 21.31.

**Table 5.** Calculated LOD, LOQ, and mean recoveries of matrix effect of analytes for spike concentrations of 10 ng mL^-1^ measured by UHPLC-MS/MS.

| No | Compound name | LOD (ng/L) | LOQ (ng/L) | ME (%) | RE (%) | R<sup>2</sup> |
| --- | --- | --- | --- | --- | --- | --- |
| 1 | Cyanidin 3,5-*O*-diglucoside | 37.93 | 125.16 | -1.42 ± 9.99 | 98.58 ± 9.99 | 0.9996 |
| 2 | Cyanidin 3-*O*-sophoroside | 15.35 | 50.65 | -9.01 ± 9.55 | 90.99 ± 9.55 | 0.9999 |
| 3 | Cyanidin 3-*O*-sambubioside | 26.02 | 85.87 | -11.91 ± 9 | 88.09 ± 9 | 0.9999 |
| 4 | Cyanidin 3-*O*-galactoside /<br>Cyanidin 3-*O*-glucoside | 28.99 | 95.65 | -7.02 ± 10.14 | 92.98 ± 10.14 | 0.9998 |
| 5 | Cyanidin 3-*O*-rutinoside | 34.71 | 114.53 | -17.26 ± 3.19 | 82.74 ± 3.19 | 0.9995 |
| 6 | Pelargonidin 3-*O*-glucoside | 3.16 | 10.41 | -5.21 ± 2.55 | 94.79 ± 2.55 | 0.9994 |
| 7 | 3-4, Dihydroxybenzoic acid | 850.34 | 2806.12 | -2.27 ± 4.26 | 97.73 ± 4.26 | 0.9993 |
| 8 | Peonidin 3-*O*-glucoside | 27.35 | 90.25 | -8.22 ± 1.84 | 91.78 ± 1.84 | 0.9996 |
| 9 | Catechin | 205.90 | 679.48 | -12.76 ± 19.09 | 87.24 ± 19.09 | 0.9966 |
| 10 | Epicatechin | 199.87 | 659.56 | -8.96 ± 6.95 | 91.04 ± 6.95 | 0.9989 |
| 11 | Caffeic acid | 319.49 | 1054.31 | 4.61 ± 5.31 | 104.61 ± 5.31 | 0.9990 |
| 12 | Delphinidin 3-*O*-rutinoside | 35.06 | 115.71 | -5.8 ± 27.02 | 94.2 ± 27.02 | 0.9983 |
| 13 | Rutin (Quercetin-3-rutinoside) | 0.13 | 0.43 | 10.54 ± 23.51 | 110.54 ± 23.51 | 0.9988 |
| 14 | Myricetin | 97.09 | 320.39 | 3.94 ± 13.21 | 96.06 ± 13.21 | 0.9937 |
| 15 | Isoquercetin/Hyperoside | 0.02 | 0.08 | 1.73 ± 2.13 | 101.73 ± 2.13 | 0.9990 |
| 16 | Kaempferol 3-*O*-rutinoside | 0.02 | 0.08 | -4.85 ± 4.73 | 95.15 ± 4.73 | 0.9995 |
| 17 | Isorhamnetin 3-rutinoside | 2.71 | 8.94 | -1.46 ± 3.76 | 98.54 ± 3.76 | 0.9988 |
| 18 | Kaempferol 3-glucoside | 0.50 | 1.65 | -1.94 ± 2.79 | 98.06 ± 2.79 | 0.9996 |
| 19 | Isorhamnetin 3-*O*-glucoside | 0.84 | 2.77 | -6.39 ± 2.37 | 93.61 ± 2.37 | 0.9984 |
| 20 | p-Coumaric acid | 150.38 | 496.24 | -10.1 ± 4.52 | 91.32 ± 4.52 | 0.9994 |
| 21 | Ferulic acid | 3.79 | 12.52 | -4.37 ± 1.24 | 95.63 ± 1.24 | 0.9999 |
| 22 | Luteolin | 60.98 | 201.22 | -7.27 ± 3.52 | 92.73 ± 3.52 | 0.9988 |
| 23 | Quercetin | 82.80 | 273.25 | 39.1 ± 21.31 | 139.1 ± 21.31 | 0.9982 |
| 24 | Naringenin | 8.82 | 29.12 | -7.41 ± 3.23 | 92.59 ± 3.23 | 0.9989 |
| 25 | Kaempferol | 11.53 | 38.05 | -9.08 ± 14.3 | 90.92 ± 14.3 | 0.9989 |
| 26 | Isorhamnetin | 3.36 | 11.08 | 19.24 ± 6.84 | 119.24 ± 6.84 | 0.9989 |

\*+ ME implies ion enhancement, – ME implies ion suppression
\*\*na = not applicable

Compounds neochlorogenic acid and chlorogenic acid exhibit inconsistent peak formation under UHPLC conditions but are detected under chromatographic conditions in an HPLC system. Chlorogenic acids (pKa = 3.5) are weak acids with carboxyl (-COOH) and phenolic hydroxyl groups (-OH) that can be deprotonated at moderately acidic pH conditions. The lower acid concentration in the aqueous mobile phase (0.01% Formic acid) of UHPLC systems might promote deprotonation of these phenolic acids. As these compounds become more anionic, their interaction with the C18 stationary phase weakens, leading to a reduction in compound retention in the column, and elution occurs earlier from the column near the solvent peak. This behavior is consistent with previous reports showing that acidified mobile phase improves retention and peak shape of phenolic acids^22,23^.

This study presents a comprehensive quantification of metabolites in American elderberry (*Sambucus nigra* subsp. *canadensis*) juices from 18 propagated accessions and three established cultivars. A streamlined and validated workflow was developed to quantify 20 bioactive metabolites using a UHPLC–MS/MS system and two additional metabolites using HPLC– MS/MS. The study also addressed matrix effects inherent to complex juice samples, which were effectively mitigated through sample dilution while maintaining detection sensitivity on the UHPLC–MS/MS platform. This approach enabled accurate and reproducible quantification of target analytes and established a robust analytical framework for metabolomic evaluation of specialty crops.

Among the metabolites analyzed, anthocyanin compounds such as cyanidin-3,5-O-diglucoside, cyanidin-3-O-sophoroside, and cyanidin-3-O-sambubioside were consistently detected across all American elderberry juices, indicating that cyanidin-based anthocyanins represent the predominant pigments in elderberry. This finding aligns with previous reports describing cyanidin derivatives as the major anthocyanins responsible for the characteristic coloration of elderberry fruits^1,5^.

The instability of myricetin observed in this study underscores the importance of considering the compound’s stability in metabolomic analysis. The stability of compounds should also be carefully evaluated during product development, as degradation of metabolites can alter both chemical composition and bioactivity of a product. Future studies should incorporate stabilization strategies, such as acidifying extraction solvents, to preserve sensitive compounds. Furthermore, despite the excellent performance of the developed analytical methods and high compound recovery, compounds such as pelargonidin 3-O-glucoside, peonidin 3-O-glucoside, catechin, epicatechin, and luteolin were not detected in any of the samples, suggesting that these compounds may be present in low concentrations. The dilution factor used in this analysis may have reduced their levels below the detection limit.

Quantification of the bioactive metabolites in juices from 21 American elderberry cultivars revealed apparent genotype-dependent variation in metabolite composition. Similar genotype-dependent differences have been reported in a previous study of European elderberry (*Sambucus nigra*)^24,25^. This finding underscores the potential for targeted breeding and cultivar selection to enhance desirable phytochemical traits, such as antioxidant capacity, color stability, and overall nutritional quality.

## 5. Conclusion

This study established and validated a high-throughput LC–MS/MS workflow for the targeted quantification of phenolic metabolites in American elderberry (Sambucus nigra subsp. canadensis) juices from 21 cultivars. Using a combination of UHPLC–MS/MS and HPLC–MS/MS platforms, 22 metabolites were successfully quantified across three major compound classes, anthocyanins, flavonoids, and phenolic acids with excellent recovery rates. The optimized UHPLC–MS/MS method demonstrated superior sensitivity, precision, and efficiency, while strategic sample dilution effectively minimized matrix effects and ensured accurate quantification in complex juice matrices.

Multivariate statistical analysis revealed apparent genotype-dependent variation between cultivars. American elderberry accessions 1896 and 2083, and the cultivar Ozark, were observed to have distinct chemical compositions, distinguishing them from other cultivars. In contrast, accession 1199 exhibited a high accumulation of cyanidin-based anthocyanins, suggesting its potential as a valuable source of natural pigment for colorant or functional food applications.

## Acknowledgments

This work is supported by the USDA-NIFA-SCRI 2021-51181-35860, “Moving American Elderberry into Mainstream Production and Processing”. We would like to acknowledge the USDA Specialty Crop Research Initiative (SCRI), the Missouri Department of Agriculture (MDA) Specialty Crop Block Grant Program (SCBGP), the Center for Agroforestry at the University of Missouri, and USDA/ARS Dale Bumpers Small Farm Research Center under agreement number 58-6020-6-001 from the USDA Agricultural Research Service for their support.

